# Chromosome-level genome assembly of the sponge *Halisarca dujardinii*

**DOI:** 10.64898/2026.02.10.705183

**Authors:** Vasiliy Zubarev, Alexander Cherkasov, Leonid Sidorov, Kim I. Adameyko, Alexander D. Finoshin, Alina Ryabova, Anastasiia Kashtanova, Guzel Gazizova, Natalia Gogoleva, Anton V. Burakov, Olga Kozlova, Elena Shagimardanova, Oleg Gusev, Kirill Mikhailov, Oksana Kravchuk, Yulia Lyupina, Ekaterina Khrameeva

**Affiliations:** Center for Molecular and Cellular Biology, Skolkovo Institute of Science and Technology, 121205 Moscow, Russia; Faculty of Bioengineering and Bioinformatics, M.V. Lomonosov Moscow State University, 119234 Moscow, Russia; Institute of Fundamental Medicine and Biology, Kazan Federal University, 420008 Kazan, Russia; Koltzov Institute of Developmental Biology, Russian Academy of Sciences, 119334 Moscow, Russia; A.N. Belozersky Institute of Physical and Chemical Biology, Lomonosov Moscow State University, 119234 Moscow, Russia; A.A. Kharkevich Institute for Information Transmission Problems, Russian Academy of Sciences, 127051 Moscow, Russia; Research Department for Limnology, Mondsee, Universität Innsbruck, 5310 Mondsee, Austria; Kazan Institute of Biochemistry and Biophysics, Federal Research Center “Kazan Scientific Center of the Russian Academy of Sciences”, 420111 Kazan, Russia; Genomics and Bioimaging Core Facility, Skolkovo Institute of Science and Technology, 121205 Moscow, Russia; Life Improvement by Future Technologies Institute, 121205 Moscow, Russia; Leibniz Institute on Aging - Fritz Lipmann Institute, 07745 Jena, Germany; Graduate School of Medicine, Juntendo University, 1138421 Tokyo, Japan

## Abstract

*Halisarca dujardinii* is a marine sponge known for its ability to completely regenerate via cell reaggregation. Here we present the first chromosome-level genome assembly of *H. dujardinii*, generated using Oxford Nanopore long reads, Illumina short reads, and Hi-C data. The final assembly spans 226.5 Mbp and is organized into 21 chromosome-scale scaffolds, representing the full nuclear genome, with an assembly N50 of 10.1 Mbp. In addition, we report the complete mitochondrial genome for *H. dujardinii*, assembled as a circular molecule and annotated to contain 14 protein-coding genes. We provide a comprehensive genome annotation comprising 14,565 nuclear protein-coding genes, of which 85.6% are functionally annotated, along with repetitive elements and non-coding RNAs. Transcript models were refined using extensive bulk and single-cell RNA sequencing data, enabling accurate annotation of 3′ untranslated regions. This new genomic resource provides a valuable foundation for investigating the molecular basis of sponge regeneration, as well as for comparative studies of sponge evolution and marine biology.

## Background & Summary

*H. dujardinii*, a member of the family *Halisarcidae*, is a widely distributed marine sponge^1^. It is best known for its extraordinary regenerative ability: the adult body of *H. dujardinii* can completely reassemble from a suspension of dissociated cells, a phenomenon known as reaggregation^2–4^. This remarkable trait makes *H. dujardinii* an attractive model for studying sponge cell biology and the fundamental mechanisms of regeneration. Research on this emerging model species is rapidly expanding. Both morphological^2,3^ and molecular^5–7^ approaches are actively used to study the reaggregation process. Recently, a whole-body single-cell RNA sequencing (scRNA-seq) dataset was generated for *H. dujardinii*^8^, allowing to identify distinct cell types and their roles during regeneration. In addition, *H. dujardinii* has served as a model for investigating diverse sponge proteins and metabolic pathways, including ferritins and iron metabolic pathways^9,10^, δ-aminolevulinic acid dehydratase^11^, neuroglobin^12^, and transglutaminase^13^. However, the absence of a high-quality genome assembly for *H. dujardinii* limits the application of omics approaches in studies involving this species.

Sponges (phylum Porifera) are among the earliest-diverging animal lineages, although their precise phylogenetic placement remains debated^14,15^. Most powerful methods in resolving large-scale phylogenies include synteny-based phylogenomic analyses that compare gene order in chromosomes across species^15^. These methods require chromosome-level genome assemblies but high-quality genomic resources for sponges are extremely limited. To date, genome assemblies are available for only 72^16^ of the 9,693^17^ described sponge species, and of these, just 8 include annotation of genes^16^. This lack of high-quality sponge genomes severely restricts comparative studies in sponge biology and marine genomics.

In this study, we present a chromosome-level genome assembly for *H. dujardinii*, generated using Oxford Nanopore long reads, Illumina short reads, and Hi-C reads. The final assembly comprises 21 chromosomes, the complete mitochondrial genome, and 201 unplaced contigs, constituting a total length of 226.5 Mbp. We also provide a high-quality genome annotation that includes a curated classification of repeats, protein-coding gene models refined with transcriptomic evidence (including scRNA-seq data for accurate 3′-UTR annotation) and accompanying functional annotations, as well as non-coding RNA genes (tRNAs and rRNAs). This assembly provides a major step toward understanding the molecular basis of *H. dujardinii*’s reaggregation ability, while also establishing a foundation for comparative analyses in sponge genomics, phylogenetics, and marine biology.

## Methods

### Sample collection

Adult specimens of *H. dujardinii* were collected from the subtidal zone of the White Sea (Russia) near the Nikolai Pertsov White Sea Biological Station (66.5533 N, 33.1039 E). Species identity was confirmed morphologically on-site. The *H. dujardinii* species is not included in the IUCN Red List or local red lists. According to local regulations, there are no specific permissions required for collecting this species in the specified location.

### Hi-C libraries preparation

To achieve chromosome-level scaffolding of the assembly, Hi-C sequencing libraries were generated according to the protocol described by Rao et al. (2014), with minor adaptations for sponge cells. Specifically, these modifications included replacing the fixation buffer with artificial seawater, adjusting reaction volumes, centrifugation speeds, and incubation times. Prior to fixation, sponge tissue fragments were rinsed with artificial seawater, minced into small pieces, and homogenized using a tissue homogenizer. The resulting cell suspension was filtered through a cell strainer, centrifuged to remove the supernatant, and flash-frozen in liquid nitrogen.

Cell fixation was carried out in a final volume of 10 mL fixation buffer (artificial seawater containing 1% formaldehyde) at room temperature with gentle rotation. The reaction was quenched by adding 2.5 M glycine to a final concentration of 0.2 M and incubated for 5 minutes at room temperature with rotation. Fixed cells were pelleted twice and resuspended in ice-cold 1× PBS.

Cells were then lysed on ice for 25 minutes using 250 μL of lysis buffer (10 mM Tris-HCl pH 8.0, 10 mM NaCl, 0.2% NP-40) supplemented with freshly added protease inhibitors. Following one wash with lysis buffer, cells were pelleted and resuspended in 0.5% SDS, then incubated for 10 minutes at 62 °C without rotation.

Chromatin digestion was performed by adding 100 U of MboI (New England Biolabs) in 250 μL of NEB2 buffer per sample, followed by overnight incubation at 37 °C with constant agitation. The resulting DNA overhangs were filled in and labeled using biotin-14-dATP and DNA Polymerase I, Large (Klenow) Fragment, along with a nucleotide mix.

After pelleting the nuclei, a ligation mix (1× NEB T4 DNA ligase buffer, 1% Triton X-100, BSA, and 400 U/μL T4 DNA Ligase) was added, and DNA fragments were ligated by overnight incubation at room temperature with gentle rotation. DNA was extracted using the phenol-chloroform method.

Samples were sheared using a Covaris LE220 instrument (fill level: 10, duty cycle: 15, PIP: 500, cycles/burst: 200, time: 58 s). Fragments of 300–500 bp were size-selected using AMPure XP beads (Beckman Coulter). Biotinylated fragments were captured using Dynabeads MyOne Streptavidin C1 beads (Thermo Fisher Scientific).

Library preparation and final amplification were conducted with the NEBNext Ultra II DNA Library Prep Kit for Illumina (New England Biolabs), following the manufacturer’s instructions. A mock PCR was performed to optimize the number of amplification cycles (eight cycles). Prior to sequencing, library size distribution was assessed using an Agilent 2100 Bioanalyzer (Agilent Technologies). The resulting in situ Hi-C libraries were sequenced using 2 × 100 bp paired-end reads on the Illumina HiSeq platform, generating 219.6 mln reads pairs and 52.90 Gb Hi-C sequencing data (Table 1).

**Table 1.** Statistics of sequencing data used for *H. dujardinii* genome assembly and annotation. *Coverage calculation based on GenomeScope2 genome size estimate, see “Technical validation” section.

| Library Type | Instrument | Layout | Total bases, GB | Coverage* | NCBI SRA Study ID |
| --- | --- | --- | --- | --- | --- |
| Illumina WGS | Illumina HiSeq 2500 | Paired-end, 249+249 | 37.00 | 186.9x | SRP519701<sup>18</sup> (present work) |
| Nanopore WGS | Oxford Nanopore MinION MK1B | Single-end | 12.84 | 64.9x | SRP519701<sup>18</sup> (present work) |
|  | Oxford Nanopore PromethION | Single-end | 16.56 | 83.6x | SRP479657<sup>19</sup> |
|  | Oxford Nanopore MinION | Single-end | 9.75 | 49.3x | SRP479657<sup>19</sup> |
| Hi-C | Illumina HiSeq 2500 | Paired-end 102+102 | 52.90 | 267.2x | SRP519701<sup>18</sup> (present work) |
| RNA sequencing | Illumina HiSeq 2500 | Single-end 57 | 50.11 | 253.1x | SRP235047<sup>20</sup> |
|  | Illumina HiSeq 2500 | Paired-end 126+126 | 73.55 | 371.5x | SRP235047<sup>20</sup> |
|  | Illumina NovaSeq 6000 | Paired-end 151+151 | 333.82 | 1,686.2x | SRP485380<sup>21</sup>; SRP558447<sup>22</sup> |
|  | Illumina HiSeq 2000 | Paired-end 101+101 | 7.43 | 37.5x | ERP013253<sup>23</sup> |
| Single-cell RNA sequencing | Illumina HiSeq 2500 | Paired-end 28 (barcode) + 91 | 54.44 | 275.0x | SRP519701<sup>18</sup> (present work) |

### Whole-genome sequencing (WGS)

To prepare libraries for Illumina sequencing, sponge tissue was first homogenized in Eppendorf plastic tubes (1.5 ml) with polypropylene pestle. Then, genomic DNA (gDNA) was extracted using a highly pure genomic DNA extraction procedure with the NucleoSpin Tissue kit (Clontech Takara), according to the manufacturer’s instructions. gDNA concentration was quantified using a Qubit 3.0 fluorometer with Quantifluor dsDNA system (Promega). Next, gDNA was fragmented using Covaris s220 (USA) DNA shearing protocol. The length of DNA fragments was estimated using Agilent Bioanalyzer 2100 (Agilent Technologies, Santa Clara, CA). Libraries for Illumina sequencing were prepared using NEBNext Ultra II DNA Library Prep Kit following the manufacturer’s protocol. The concentration of libraries was measured by the Qubit 3.0 fluorometer. The quality and length of libraries was measured on Agilent 2100 Bioanalyzer using DNA High Sensitivity chip (Agilent Technologies, Santa Clara, CA). The library was sequenced on the Illumina HiSeq 2500 instrument in 249+249 bp mode, generating 37.00 Gb of data and 186.9x genomic coverage in total (Table 1).

To prepare libraries for Oxford Nanopore (ONP) sequencing, frozen sponge tissue was first ground under liquid nitrogen using a ceramic pestle and mortar. Then, genomic high-molecular-weight (HMW) DNA was extracted using the phenol-chloroform method according to standard protocols. Additional purification was performed using CTAB-mediated DNA precipitation^24^. The resulting HMW DNA was treated with the Short Read Eliminator Kit (XS) (Circulomics) to remove fragments shorter than 10 kb. The Ligation Sequencing Kit (SQK-LSK109, Oxford Nanopore Technologies, UK) was employed to prepare Nanopore sequencing libraries from total DNA without shearing. The manufacturer’s protocol was used with the following modifications: NEB end-prep and repair times were extended sixfold, to 30 min at 20°C and 30 min at 65°C, and adapter ligation time was extended to 1 h. The libraries were sequenced on the MinION MK1B (Oxford Nanopore Technologies, UK) using flow cell FLO-MIN106D R9.4.1 for Nanopore sequencing. The MinKNOW software package (v.19.12.5) was used for basecalling and demultiplexing reads. Sequencing was conducted three times, and the resulting 12.84 Gb of data were combined, generating 64.9x genomic coverage in total (Table 1). N50 of obtained ONP reads is 13,241 bp and mean length is 6,236.8 bp.

### Contig-level *de novo* genome assembly

Raw Illumina and Oxford Nanopore genomic sequencing data^18^ were assessed for quality using FastQC v.0.11.8 prior to genome assembly. Illumina libraries were trimmed with fastp v.0.23.2^25^ (--detect_adapter_for_pe --cut_tail --cut_front --trim_poly_g --cut_mean_quality 20 --length_required 100 --average_qual 25). In addition to our own ONP library, we downloaded *H. dujardinii* ONP data from NCBI SRA^19^ (see Table 1) and merged it with our dataset, resulting in a total ONP data volume of 39.15 Gb. ONP reads were not filtered or trimmed prior to assembly because the majority of assemblers have internal cutoffs for Oxford Nanopore read length and quality or perform read correction as part of the assembly workflow. To select an optimal strategy for draft contig-level genome assembly, we evaluated several *de novo* assemblers: wtdbg2 v.2.5^26^, flye v.2.8.1^27^, hifiasm v.0.25.0^28^, NextDenovo v.2.5.2^29^ (Figure 1, Table 2). Previous studies on mollusc genomes have demonstrated that assemblies generated from the longest ONP reads providing approximately 60x genome coverage can be more accurate than those using the complete read set^30^. Therefore, long-read assemblers Flye, hifiasm, and NextDenovo were run using two long-read input strategies: (i) all available ONP reads (39.15 GB, N50 = 14,833 bp) and (ii) a subset corresponding to 60x genome coverage of the longest reads (11.9 GB, N50 = 30,109 bp). Assembly quality was evaluated using total assembly length, contig N50 (calculated with seqkit v.2.10.1^31^), and complete Best Universal Single-Copy Orthologs (BUSCO) score (calculated using compleasm v.0.2.7^32^ with the metazoa_odb12^33^ BUSCO lineage dataset). To avoid underestimation of BUSCO completeness due to low single-nucleotide quality in assemblies generated exclusively from ONP reads, all assemblies were polished prior to BUSCO analysis using pilon v.1.24^34^ (--fix “snps,indels” --diploid --changes). Polishing was performed using our PE250 Illumina genomic library^18^, trimmed as described above and aligned to the corresponding assemblies with bwa v.0.7.17^35^ (bwa mem-T 30).

**Table 2.**
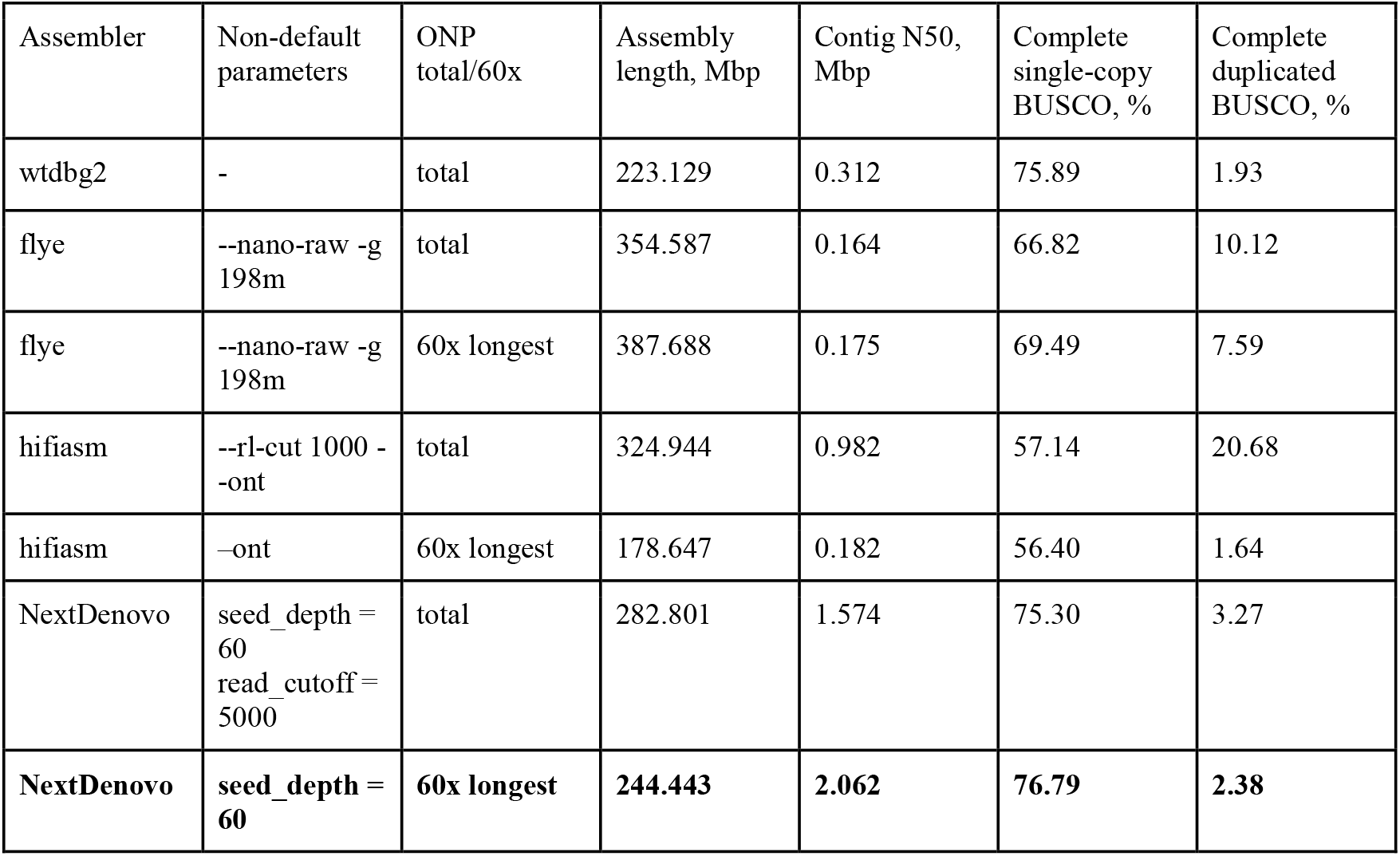
Summary of draft contig-level genome assemblies. The selected assembly variant is highlighted in bold.

**Figure 1.**
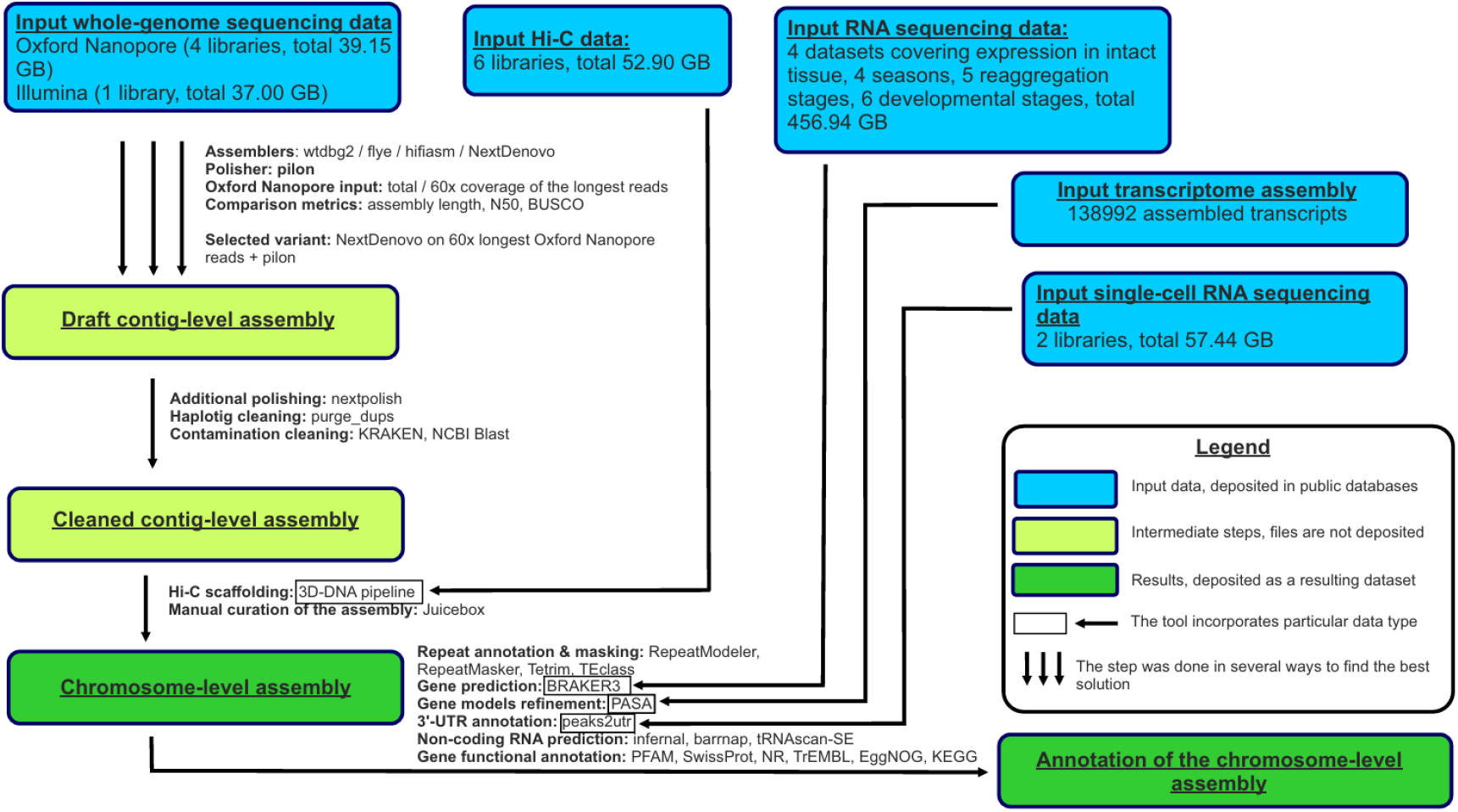
Workflow diagram illustrating the major steps of this study.

The draft contig-level assembly selected for downstream analyses was the NextDenovo assembly generated using the longest ONP reads corresponding to approximately 60x genome coverage. This assembly simultaneously exhibited a high contig N50 and BUSCO scores, while its total length was reasonably close to the predicted genome size (198 Mbp; see Technical Validation section). However, the assembly size exceeded the expected value, suggesting the presence of redundant haplotypic contigs (haplotigs).To remove putative haplotigs, the draft assembly was processed with purge_dups v.1.2.5^36^. As subsequent steps (e.g., RNA-seq read mapping) revealed insufficient single-nucleotide quality, we performed additional polishing with NextPolish^37^. Bacterial contamination has previously been reported in genome assemblies of other sponges^38,39^. To identify and remove contaminant sequences, contigs from the draft assembly were taxonomically classified using KRAKEN v.2.0.8^40^. Contigs assigned to bacteria were further validated using the NCBI BLASTN online tool. All sequences that were classified as bacterial by Kraken and had NCBI BLASTN best hits corresponding to bacteria were removed. The resulting assembly was then screened for residual adapter and foreign sequences using the NCBI Foreign Contamination Screen^41^. Collectively, haplotig removal and contamination filtering reduced the total assembly size from 244.4 Mbp (in draft contig-level assembly) to 226.5 Mbp (in cleaned contig-level assembly).

### Chromosome-level scaffolding and validation via metaphase imaging

To generate a chromosome-level assembly, the cleaned contig-level assembly was scaffolded using the 3D-DNA pipeline v.180922^42^, with parameters tuned to detect misjoins at large resolutions (--mapq 30 --editor-fine-resolution 1000 --editor-coarse-resolution 50000 --editor-coarse-region 250000 --editor-repeat-coverage 40 --mode diploid). The obtained chromosome-level assembly was then polished by manual curation to improve its accuracy using Juicebox v.2.17.00. The Hi-C map for the curated chromosome-level assembly was prepared using distiller-nf v.0.3.4^43^ (Figure 2a). This Hi-C-based scaffolding procedure dramatically improved the assembly contiguity, increasing the scaffold N50 to 10,133 kbp (from 2,062 kbp at the contig stage) and anchoring 96.54% of the assembled bases into 21 scaffolds (Table 3). The resulting scaffolds are presumed to represent the 21 chromosomes of *H. dujardinii*, with the total length of 218.7 Mbp and 201 unplaced contigs.

**Table 3.** Summary statistics of the *H. dujardinii* chromosome-level genome assembly.

|  |  |
| --- | --- |
| Total length | 226,506,016 bp |
| Length of the chromosome part | 218,679,722 bp |
| Number of chromosomes | 21 |
| Number of unplaced sequences | 201 |
| N50 | 10,133,527 bp |
| Number of protein-coding genes | 14,565 nuclear, 14 mitochondrial |
| Number of proteins (including isoforms) | 19,581 nuclear, 14 mitochondrial |
| Compleasm score (complete genes) | 86.60% |

**Figure 2.**
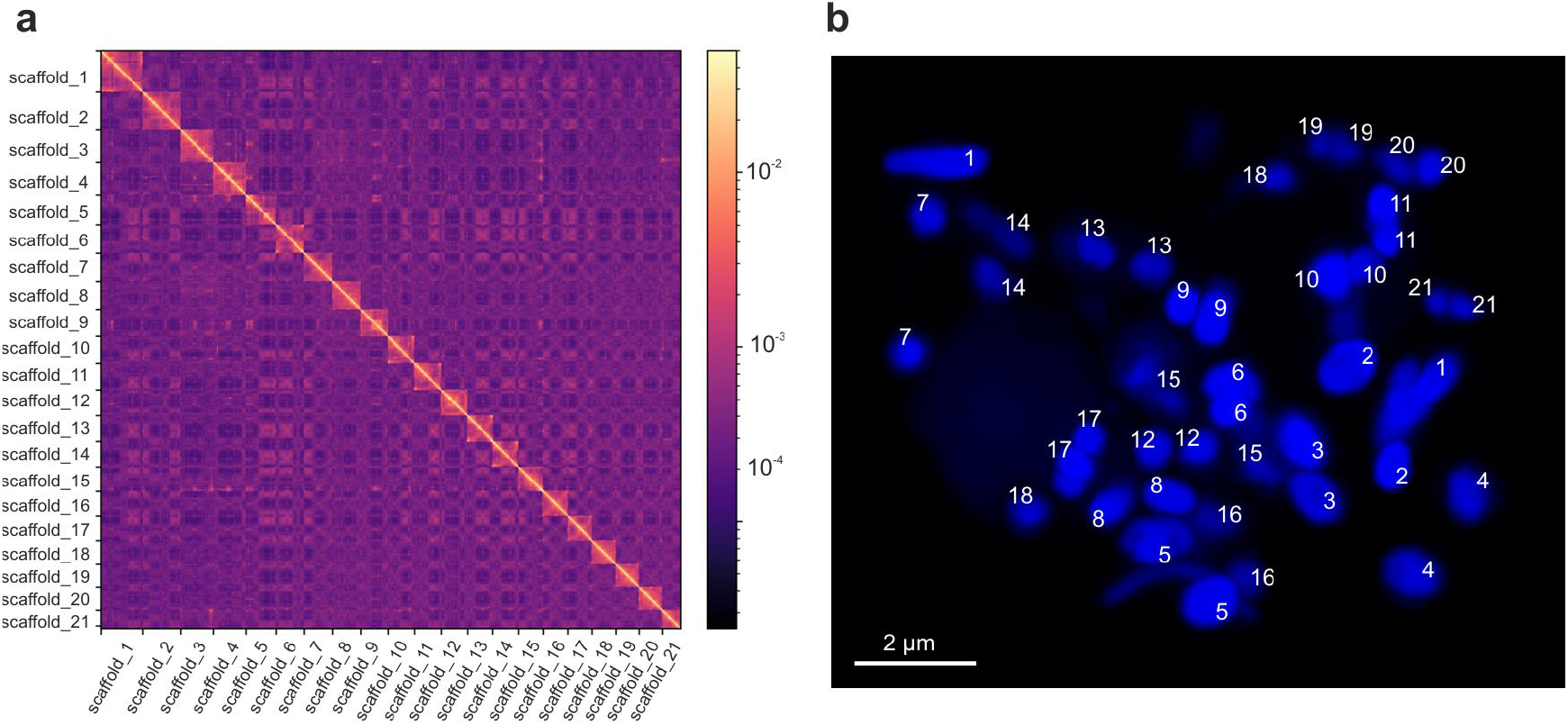
Chromosome-level assembly of the *H. dujardinii* genome. (**a**) Hi-C contact map at 100 kbp resolution, showing interaction frequencies used to scaffold contigs into the chromosome-level assembly. Unplaced contigs are not shown. (**b**) Confocal image of mitotic metaphase chromosomes prepared from *H. dujardinii* cells, stained with Hoechst-33342 to visualize individual chromosomes. Scale bar, 2 µm.

To experimentally validate the number of chromosomes in *H. dujardinii*, we analyzed chromosomes at the metaphase stage of mitosis, when they are maximally condensed and therefore most visible. To capture confocal images of chromosomes, the sponge was first incubated in filtered seawater (FSW) containing 0.007% colchicine for 12 hours at 6–8 °C (1). Subsequently, it was mechanically dissociated into individual cells as previously described (2). Dissociated cells were incubated for 1 h in FSW in six-well plates with a coverslip at the bottom to mount cells, and then the mounted cells were treated with a hypotonic solution of 0.4% KCl for 30 minutes. After hypotonic treatment, the solution was discarded, and cells were fixed by washing twice in a freshly prepared cold fixative solution (3:1 methanol:acetic acid), with each wash lasting at least 30 minutes. Coverslips were then air-dried and rinsed with 50% acetic acid.

Next, coverslips were washed in PBS for 10 minutes and stained with Hoechst-33342 (Invitrogen/Thermo Fisher Scientific, Waltham, MA, USA) for 5 minutes to visualize individual chromosomes. Finally, coverslips were washed in PBS for 10 minutes and mounted using antifade liquid (Amerigo Scientific, NY, USA) on slides for confocal imaging using the Carl Zeiss LSM 880 microscope (Carl Zeiss AG, Oberkochen, Germany) and the manufacturer’s software Zen Blue. The resulting images of *H. dujardinii* chromosomes confirm that the number of scaffolds obtained in our chromosome-level assembly matches the number observed in the images of metaphase chromosomes – 21 chromosomes (Figure 2b).

### Repeat annotation

Repetitive elements in the assembled *H. dujardinii* genome were annotated and masked using RepeatModeler2 and RepeatMasker v.4.1.7-p1^44^. To obtain a comprehensive set of sponge repeat families, we applied RepeatModeler2 to a database comprising all RefSeq sponge genome assemblies, including the newly assembled *H. dujardinii* genome. Thus, this database included seven species: *Amphimedon queenslandica*^45^, *Corticium candelabrum*^46^, *Dysidea avara*^47^, *Halichondria panicea*^48^, *Oscarella lobularis*^49^, *Sycon ciliatum*^50^, *and H. dujardinii*. This analysis produced 7,251 consensus repeat sequences, which were then processed using TEtrimmer v.1.4.0^51^ with default parameters. The curated repeat library consisted of 612 refined sequences, of which 303 were annotated and 309 remained unclassified. To annotate the remaining set of unclassified repeat sequences, we further applied the TEclass2^52^ tool with a probability threshold of 0.7 for correct classification. This approach resulted in successful classification of an additional 209 out of 309 previously unclassified sequences.

This repeat library was then used for soft masking with RepeatMasker^44^ through a three-step procedure. First, we masked only simple and low-copy repeats. Next, annotated complex repeats were masked in the output genome from the previous step. Finally, we masked unclassified complex repeats in the same manner. This iterative masking approach allowed us to prioritize the annotation of elements with homology to known repeat families, minimizing inadvertent matching to unknown repeats. Custom scripts provided in the GenomeAnnotation GitHub repository (https://github.com/darencard/GenomeAnnotation) were used to facilitate this repeat masking and annotation workflow. As a result, 47.29% of the assembled *H. dujardinii* genome was annotated as repetitive and masked accordingly (Figure 3a), including long interspersed nuclear elements (LINEs; 11.30%), DNA transposons (7.23%), and long terminal repeat elements (LTRs; 5.14%) as the major identified repeat classes.

**Figure 3.**
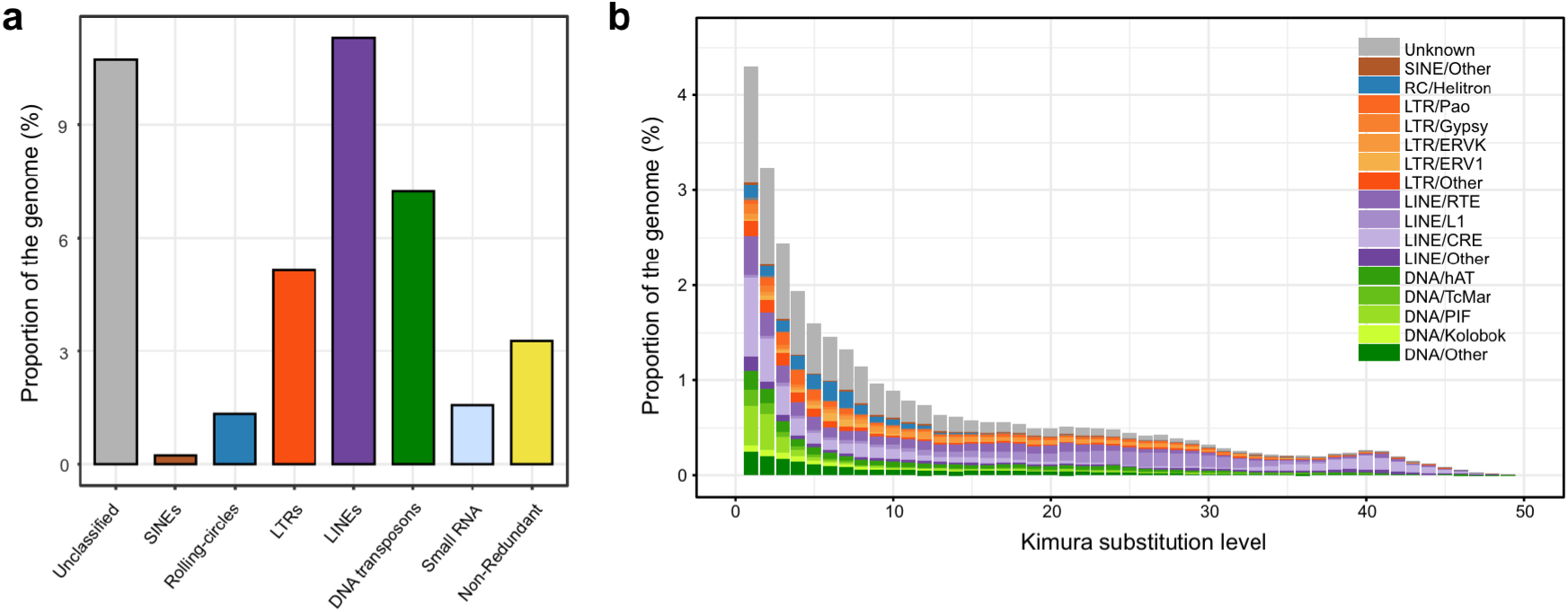
Repetitive elements in the assembled *H. dujardinii* genome. (**a**) Breakdown of repeat classes in the assembled *H. dujardinii* genome, with proportions for major classes including long interspersed nuclear elements (LINEs), long terminal repeat elements (LTRs), DNA transposons, and others. (**b**) Evolutionary landscape of repeat families, showing their genomic abundance across Kimura substitution levels. The Kimura distance estimates the number of nucleotide substitutions per site between each transposable element (TE) copy and its family’s consensus sequence, correcting for differing rates of transitions and transversions. This divergence serves as a proxy for the time since insertion: lower Kimura values reflect more recent TE activity, while higher values indicate older insertions that have accumulated more mutations over time.

Finally, we studied the evolutionary history and dynamics of annotated mobile genetic elements. We visualized the proportion of the genome covered by transposable element (TE) families across different levels of sequence divergence using RepeatMasker’s *createRepeatLandscape*.*pl* script. This analysis uncovered a recent proliferation of previously uncharacterized mobile elements, along with two known groups: Kolobok elements, a class of DNA transposons characterized by a DDE transposase domain and often associated with terminal inverted repeats, and Crypton-related elements (CRE), which are tyrosine recombinase-containing DNA transposons often found in lower eukaryotes^53^ (Figure 3b). Such a profile, dominated by low-divergence peaks, suggests ongoing TE activity in *H. dujardinii*, with continual insertion of young elements that have yet to accumulate many mutations.

### RNA-Seq Data Preparation

To capture both life-stage-specific and housekeeping transcripts for comprehensive annotation, we collected bulk and single-cell RNA-Seq data of *H. dujardinii* from four independent sources representing diverse biological conditions:

- Institute of Developmental Biology RAS (IDB). We used six paired-end (126+126 bp) and 27 single-end (57 bp) bulk RNA-Seq libraries from intact body, dissociated cells, and cell aggregates 24 hours post dissociation of *H. dujardinii* collected during all four seasons (NCBI SRA accession SRP235047^20^). Raw reads were trimmed and quality-checked with FastQC v.0.12.1^54^, MultiQC v.1.28^55^ and fastp v.0.24.0^25^. We generated six de novo transcriptome assemblies from paired-end reads using Trinity v2.8.5^56^, clustering them subsequently using CD-HIT-EST v.4.8.1^57^ with parameters “-c 0.98-aS 0.95-aL 0.90”. Additionally, paired-end reads were aligned to the genome with STAR v2.7.11b in two passes to maximize splice junction detection. Then, we constructed genome-guided transcript assemblies with StringTie v3.0.0. Single-cell reads from NCBI SRA accession SRP519701^18^ (SRR29818416-SRR29818419) were aligned to the genome assembly using CellRanger v.9.0.1 (10X Genomics) and used for subsequent 3’UTR annotation.
- Australian National University (ANU). We used the published Trinity assembly of *H. dujardinii* laboratory-grown, contamination-free juveniles (NCBI TSA ID: HADA01^58^). We also performed a new genome-guided StringTie assembly on this dataset (NCBI SRA accession ERP013253^23^).
- Saint Petersburg State University (SPbU). We performed genome-guided StringTie assemblies using six RNA-Seq libraries (NCBI SRA accessions SRR27694415, SRR27694418, SRR27694419, SRR27694421, SRR27694423, SRR27694424^21^) covering key developmental stages: early cleavage, middle cleavage, smooth prelarva, folded prelarva, disphaerula, and free-swimming larva.
- Lomonosov Moscow State University (MSU). We performed genome-guided StringTie assemblies using six RNA-Seq libraries (NCBI SRA accessions SRR32050810, SRR32050814, SRR32050818, SRR32050821, SRR32050824, SRR32050825) covering key stages of sponge reaggregation: intact tissues, primary multicellular aggregates (6 hours post-dissociation, hpd), early-stage primmorphs (24 hpd), true primmorphs (72 hpd), primmorphs with a developing aquiferous system, and fully reconstructed sponges.

### Gene structure prediction

To generate gene models for *H. dujardinii*, we applied the combined RNA-seq and homology-based annotation pipeline BRAKER v.3.0.8^32,59–73^, using pooled RNA-seq alignments from the Institute of Developmental Biology (IDB)^20^ and Australian National University (ANU)^23^ datasets. Reads were first filtered with fastp v.0.24.0^25^, aligned to the genome with STAR v.2.7.11b^74^, and low-confidence alignments (MAPQ < 30) were removed using samtools v1.21^75^. The resulting BAM files were supplied to BRAKER3 as RNA evidence for gene prediction. Splice junctions and exon-intron boundaries were manually validated in the Integrative Genomics Viewer (IGV) v2.16.0^76^. Protein homology evidence for BRAKER3 was provided by the *metazoa_odb12* dataset from OrthoDB v.12^33^ and proteomes of the six above-mentioned sponge species from the NCBI RefSeq database^45–50^.

To incorporate untranslated regions (UTRs) and refine transcript structures, we used PASA v.2.5.3^77,78^ with transcript sets both from *de novo* IDB and ANU Trinity transcriptomes and merged StringTie assemblies from all four RNA-seq sources - IDB, ANU, SPbU and MSU^20–23^) using gmap and minimap2 aligners. We performed three iterative rounds of PASA’s *--annot_compare* reannotation module. To filter out erroneous long intron alignments, we set *--MAX_INTRON_LENGTH 30000*.

The 3′UTRs were further refined using IDB’s scRNA-seq data^18^ enriched for transcript 3′ ends. These reads were aligned using CellRanger v.9.0.1^79^, and 3′ end peaks were used to guide UTR refinement via peaks2utr v.1.4.1^80^. This UTR refinement was specifically important for accurate mapping of scRNA-seq reads (see below).

In total, 14,565 gene models were predicted, comprising 19,580 transcripts after the PASA and peaks2utr reannotation (on average, 1.34 isoforms per gene). Across the genome, the average gene density is approximately 6.53 genes per 100-kb window; windows without genes are rare (2.86%), and the median intergenic distance is 8,780 bp (Figure 4a). We selected the longest isoform as the canonical transcript for each gene. Among these 14,565 canonical transcripts, 3,124 (21.4%) include an annotated 5′ UTR and 10,657 (73.2%) include an annotated 3′ UTR. Monoexonic transcripts account for 4,639 (31.9%) of canonical isoforms, nearly half of which (n = 2,185) lack UTR annotations; the remaining 9,926 (68.1%) are multiexonic, with an average of seven exons per transcript (Figure 4b). The median length of monoexonic transcripts is 1,005 bp (597 bp without annotated UTRs versus 1,371 bp with UTRs), whereas multiexonic transcripts have median lengths of 7,501 bp pre-splicing and 1,755 bp post-splicing (Figure 4c). Coding exons in multiexonic genes are short (median 117 bp), while exons spanning UTRs tend to be longer (Figure 4d). Coding-region introns display a median length of 517 bp, with a secondary peak near 70 bp, reflecting two distinct intron size classes (Figure 4e). UTR regions are rarely intronated, with an average of 1.2 exons per 5′ UTR and 1.0 exon per 3′ UTR, and account for fewer than 1.5% of all introns (n = 1,165 out of 84,876). The mean GC content is 43.91%, and regions of elevated GC content generally coincide with gene-rich windows (Figure 5).

**Figure 4.**
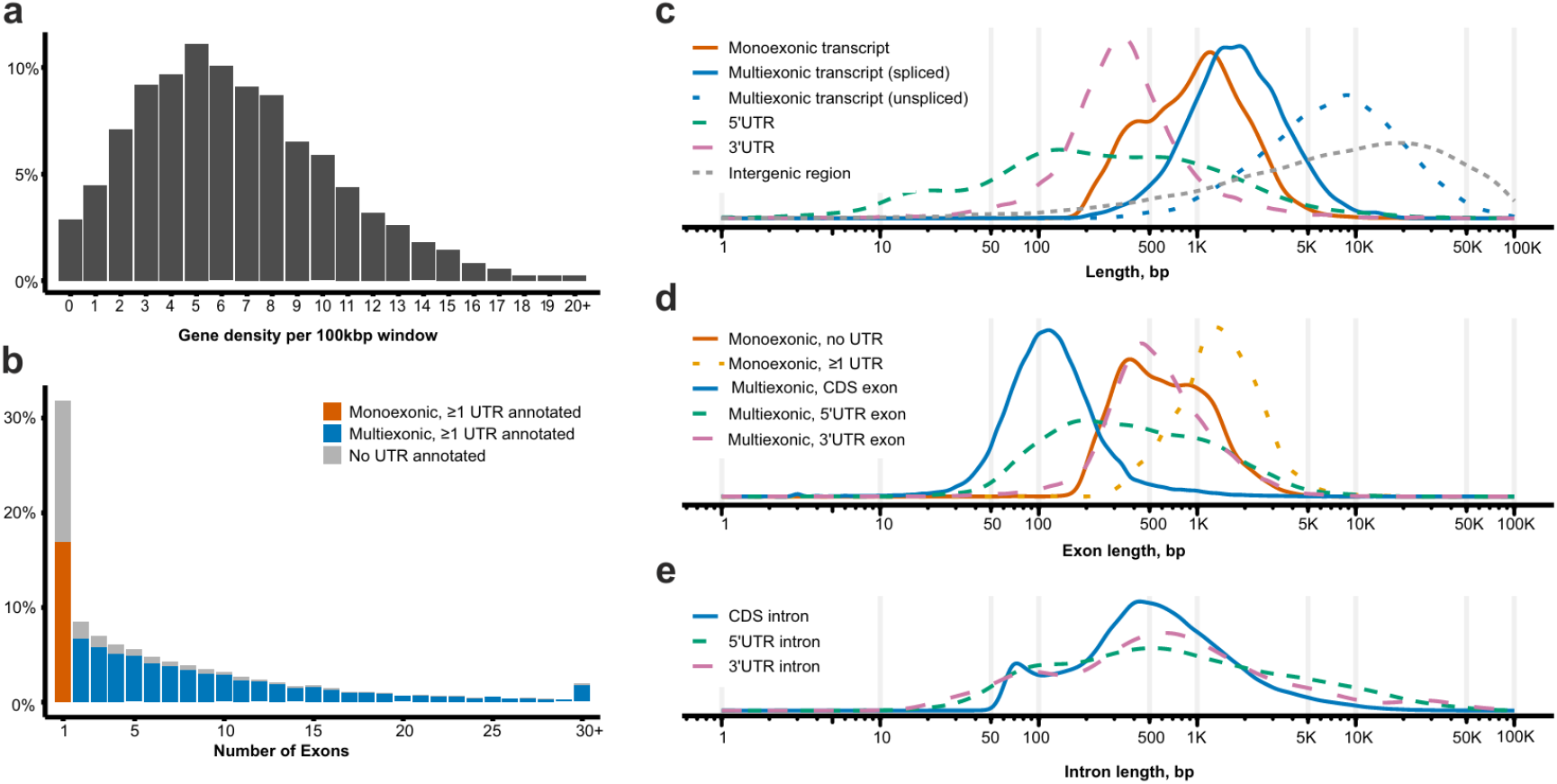
Summary of protein-coding genes’ features in *H. dujardinii*. (**a**) Gene density per 100 kbp non-overlapping genomic regions. (**b**) Exon count distribution for protein-coding transcripts. (**c**) Distributions of transcript elements’ lengths and intergenic distances. (**d**) Exon length distributions. (**e**) Intron length distributions.

**Figure 5.**
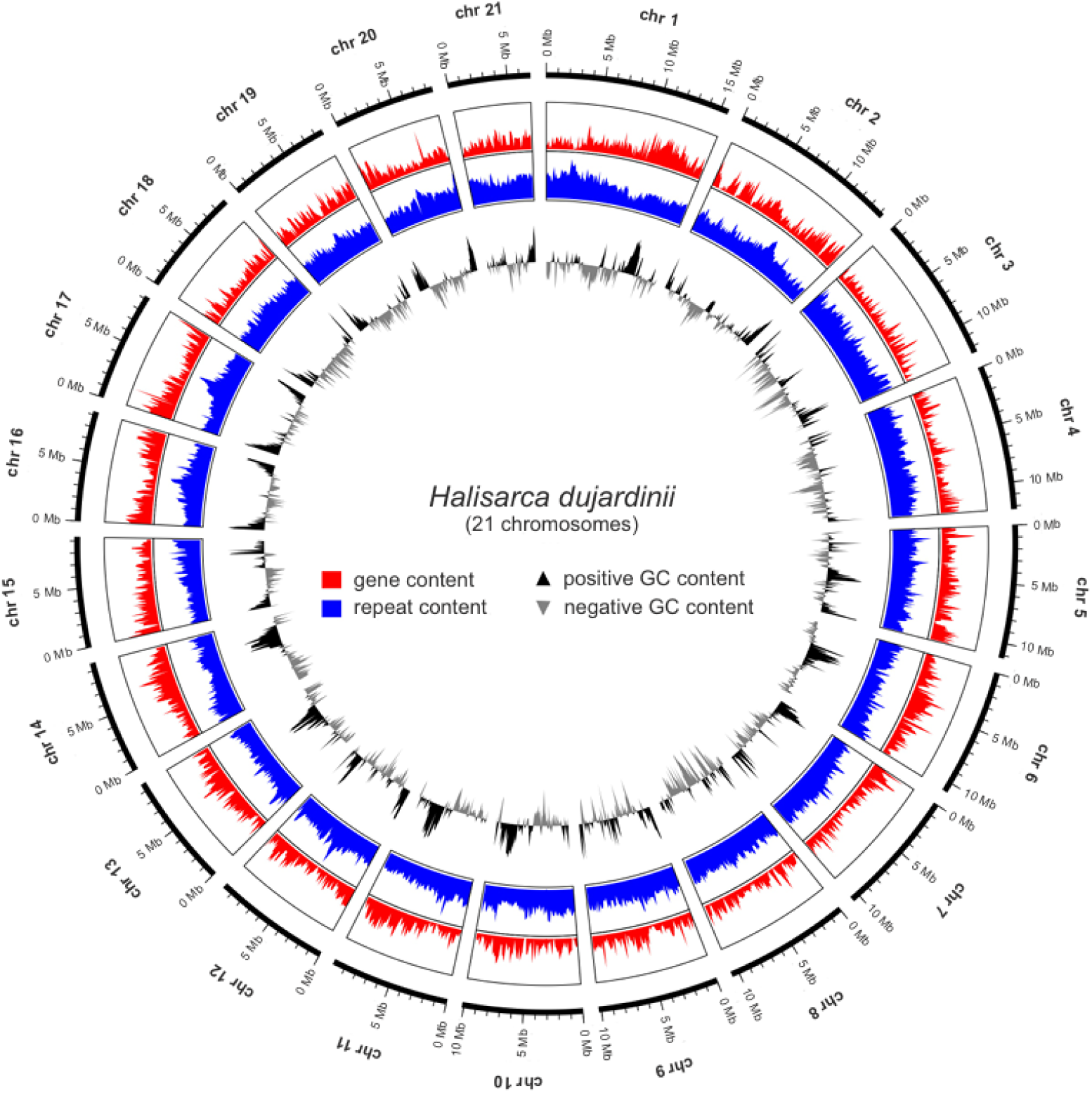
Circos plot of *H. dujardinii* chromosomes. A circular representation showing, from outer to inner rings: chromosome ideograms, gene density (number of genes per window), repeat density (percentage of bases masked per window), and GC content profile (percentage of GC bases per window).

Further, tRNA and rRNA genes were predicted using tRNAscan-SE v.2.0.12^81^ and barrnap v.0.9 (https://github.com/tseemann/barrnap), respectively. Of note, rRNA genes cluster in the nucleolar organizing region located on chromosome 7.

### Functional annotation of genes

The functional annotation of *H. dujardinii* genes was performed using the Trinotate v.4.0.2 pipeline^82^. The annotation process included conducting BLASTP and BLASTX searches (implemented in DIAMOND v.2.1.11^65^) against the SwissProt database; searching Pfam domains using HMMer v.3.4^83^; assigning Kyoto Encyclopedia of Genes and Genomes (KEGG) categories with GHOSTX in KEGG Automatic Annotation Server^84^, and inferring functions by orthology using EggNOG-mapper v.2.1.12^85,86^ (Figure 6a). Additionally, the homology search was conducted against the TrEMBL database using mmseqs2^87^. Signal peptides were predicted using SignalP v.6.0^88^, and transmembrane domains were predicted using TMHMM v.2.0^89^ (Figure 6b).

**Figure 6.**
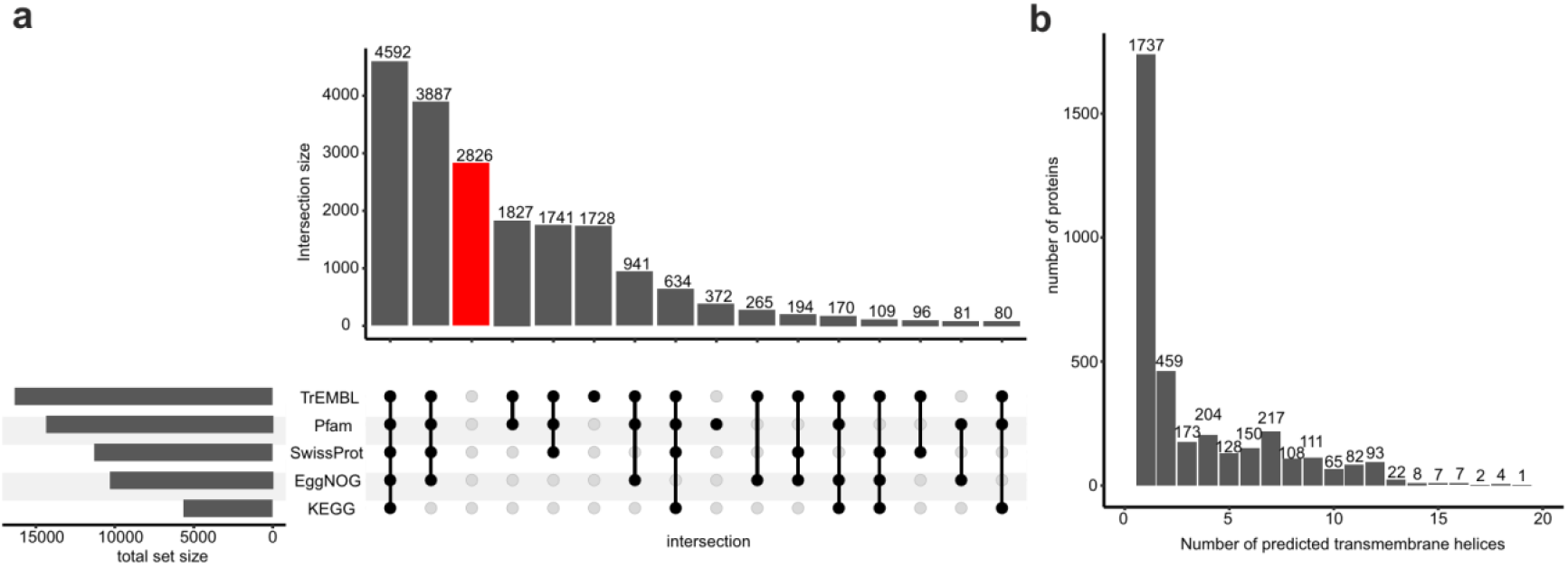
Functional annotation of genes. (**a**) UpSet plot illustrating overlaps among transcript sets annotated using five complementary approaches: (1) BLASTP search against the TrEMBL database, (2) protein domain identification via HMM search against Pfam, (3) BLASTP and BLASTX searches against the SwissProt database, (4) orthology inference using EggNOG-mapper, (5) search against the KEGG orthology database. Intersections comprising fewer than 50 transcripts are not shown. Transcripts lacking functional annotation are highlighted in red. (**b**) Distribution of proteins by the number of predicted transmembrane domains.

In total, 4,592 proteins out of 19,581 (23.5%) were annotated by all these methods, and 16,755 were annotated by at least one method (85.6%). Signal peptides were predicted for 1,973 proteins, and 3,585 proteins were shown to contain transmembrane domains. The distribution of transmembrane domain counts displays a pronounced peak at membrane-attached proteins with a single transmembrane domain, followed by a long tail of transmembrane proteins with increasingly complex topologies, containing up to 19 transmembrane domains (Figure 6b).

### Mitochondrial genome assembly and annotation

For certain omics analyses, it is crucial to include a high-quality mitochondrial genome in the reference – for example, in scRNA-seq analysis, where the fraction of mitochondrial gene expression serves as a quality control metric. Therefore, we made an effort to include the mitochondrial genome in our assembly. We assembled the mitochondrial genome of *H. dujardinii* from whole-genome Illumina sequencing data from Australian University^23^. We used the *get_organelle_from_reads*.*py* script from the GetOrganelle v.1.7.7.1 package^90^. This produced a circular genome of length 19,277 bp. We annotated this genome using the DeGeCI v.1.1 online server^91^ and MITOS v.2.1.7^92^ via the Proksee online server^93^. This annotation identified 14 protein-coding genes, 25 tRNA genes, and 2 rRNA genes.

To summarize, the new chromosome-level genome assembly of *H. dujardinii* includes 21 chromosomes, mitochondrion, and 201 unplaced contigs (Table 3). The assembly N50 is 10,133,527 bp. The combined length of the assembled chromosomes in our assembly is 218,679,722 bp.

### Data Records

Our chromosome-level genome assembly is available at NCBI GenBank database^94^.

Our whole-genome sequencing data for *H. dujardinii*, including Illumina and Oxford Nanopore reads, have been deposited in the NCBI SRA database^18^, accessions: SRR31896582 (Illumina), SRR31896581 (Oxford Nanopore).

Our Hi-C data for *H. dujardinii* have been deposited in the NCBI SRA^18^, accessions: SRR33832908-SRR33832915.

Annotation (GFF) files and a table detailing functional annotation are available at Zenodo^95^.

### Technical Validation

To assess completeness of the chromosome-level assembly^94^, we applied BUSCO^59^ method implemented in the compleasm package^32^. We compared our assembly to all six sponge genomes available at RefSeq (*Amphimedon queenslandica*^45^, *Corticium candelabrum*^46^, *Dysidea avara*^47^, *Halichondria panicea*^48^, *Oscarella lobularis*^49^, *Sycon ciliatum*^50^) in both genome and protein modes against the metazoa_odb12 BUSCO dataset from OrthoDB v.12^33^. According to the genome mode analysis, our *H. dujardinii* assembly contains 77.8% complete BUSCO genes. This score is comparable to six available RefSeq sponge assemblies, which contain from 76.3% (*Sycon ciliatum*) to 87.2% (*Corticum candelabrum*) complete BUSCO genes (Figure 7a, left). For the protein mode analysis, we extracted amino acid sequences by coordinates using the *agat_sp_extract_sequences*.*pl* script from the GFF file manipulation package AGAT^96^. According to the protein mode analysis, gene annotation for our *H. dujardinii* assembly contains 86.6% complete BUSCO genes. This score is also comparable to six RefSeq sponge annotations that contain from 86.2% (*Sycon ciliatum*) to 90.3% (*Dysidea avara*) complete BUSCO genes (Figure 7a, right).

**Figure 7.**
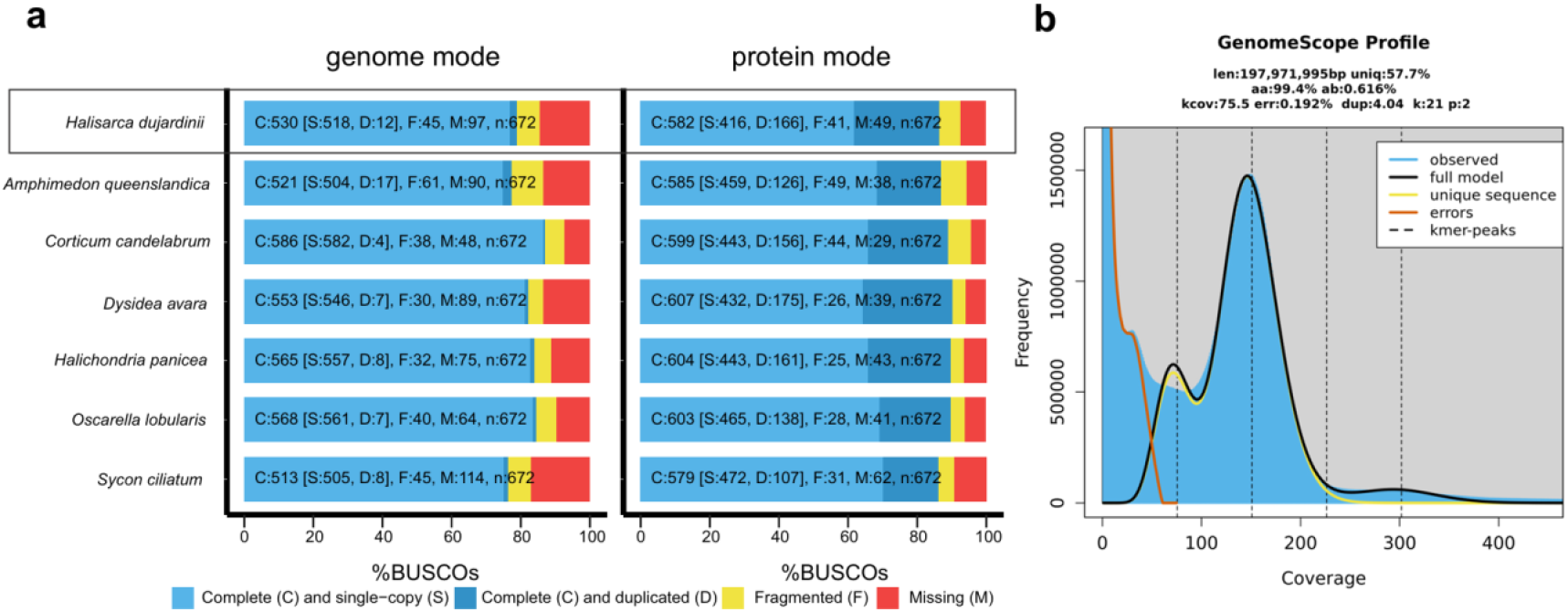
Technical validation of the novel *H. dujardinii* assembly. (**a**) BUSCO completeness scores for our novel *H. dujardinii* assembly and six sponge genomes available at RefSeq, shown separately for genome and protein modes. (**b**) K-mer frequency spectrum of *H. dujardinii* whole-genome sequencing data, overlaid with the fitted GenomeScope2 model.

To validate the genome size of our assembly, we employed an independent k-mer-based method. We generated a k-mer frequency spectrum with Jellyfish v.2.2.10^97^ and fitted it using GenomeScope2 v.2.0.1^98,99^. The model estimates the genome size of 198 Mbp (Figure 6b), indicating that our chromosome-level assembly size is 114.5% of the estimated genome length and the total length of chromosomes excluding unplaced sequences is 110.5% of the estimated genome length. The proportion of repetitive sequences masked by RepeatMasker in our assembly (47.29% of the genome length) closely matches the GenomeScope2 prediction (42.3% of repetitive sequences).

To evaluate how the 3′-UTR refinement procedure (see above) improves scRNA-seq mapping accuracy, we compared CellRanger^79^ metrics before and after running peaks2utr (Table 4). As expected, the overall read mapping rate to the genome remained unchanged. However, a subset of reads previously assigned to intergenic regions were shifted to exons. This improved both the total count of detected cells and the median number of genes per cell.

**Table 4.** CellRanger^79^ mapping and cell-counting metrics for our *H. dujardinii* assembly before and after the 3’-UTR refinement procedure. The metrics are listed for two replicates from the IDB scRNA-seq dataset^18^.

| Metric | Before 3'-UTR annotation |  | After 3'-UTR annotation |  |
| --- | --- | --- | --- | --- |
|  | Replicate 1 | Replicate 2 | Replicate 1 | Replicate 2 |
| Reads mapped to genome, % | 87.7 | 81.4 | 87.7 | 81.4 |
| Reads Mapped Confidently to Intergenic Regions, % | 23.1 | 32.6 | 10.6 | 9.7 |
| Reads Mapped Confidently to Intronic Regions, % | 4.3 | 4.8 | 4.3 | 4.7 |
| Reads Mapped Confidently to Exonic Regions, % | 43.5 | 31.5 | 57.0 | 55.2 |
| Reads Mapped Confidently to Transcriptome, % | 45.3 | 33.2 | 58.8 | 57.0 |
| Estimated number of cells | 10,406 | 12,252 | 10,840 | 13,704 |
| Median genes per cell | 1,489 | 1,335 | 1,826 | 1,929 |

To compare our chromosome-level genome assembly and annotation with the previously published scaffold-level version^19,100,101^, we assessed BUSCO completeness and evaluated mapping performance for bulk RNA-seq and single-cell RNA-seq (scRNA-seq) data. As an independent validation dataset for bulk RNA-seq, we used 22 RNA-seq libraries from the NCBI SRA accession SRP485609^102^ (not used at any step annotation of genome annotation). These RNA-seq libraries were filtered using fastp v.0.24.0^25^, randomly downsampled to one million read pairs with seqkit v.2.9.0^31^, and mapped to the annotated genome using STAR v.2.7.11b^74^. Available scRNA-seq data for *H. dujardinii* are limited; therefore, the same two scRNA-seq libraries^18^ were used for testing and for annotation. These libraries were mapped to the annotated genome using CellRanger v.9.0.1^79^. Multiqc v.1.28^55^ was used to aggregate mapping statistics and quality metrics from all analyses.

The results (Table 5) show that, in addition to a dramatically increased N50 value, our assembly achieves higher BUSCO completeness. Mapping rates further indicate that our version provides a slightly better reference for bulk RNA-seq read mapping and a pronounced enhancement for scRNA-seq read mapping, mostly due to more accurate 3’-UTR annotations. In addition, unlike the previously published assembly^100,101^, our version includes a fully assembled and annotated circular mitochondrial genome.

**Table 5.** Comparison of our chromosome-level *H. dujardinii* genome assembly with the previously published draft version^100,101^.

| Metric | Our chromosome-level assembly<sup>94</sup> | Draft assembly<sup>100,101</sup> |
| --- | --- | --- |
| N50, kbp | 10,133.5 | 785 |
| Number of genes | 14,565 | 13,989 |
| Complete BUSCO genes (genome mode), % | 78.87 | 76.04 |
| Complete BUSCO genes (protein mode), % | 86.60 | 85.56 |
| RNA-seq mapping rate (unique mappings), average of 22 libraries, % | 84.31 | 79.03 |
| RNA-seq mapping rate (unique mappings assigned to genes), average of 22 libraries, % | 68.60 | 65.93 |
| scRNA-seq reads confidently mapped to exonic regions, average of two libraries, % | 56.1 | 41.7 |
| Mitochondrial genome, kbp | 19.3 | NA |

### Usage Notes

- FASTA and GTF files were validated for compatibility with the reference making modules of STAR v.2.7.11, HISAT2 v.2.2.1 and CellRanger v.9.0.1.
- In the provided GFF and GTF annotation files, feature identifiers are prefixed to denote the major annotation category. The following prefixes are used: “SkHd3_prot_” for nuclear protein-coding genes, “SkHd3_MT_” for mitochondrial genes, “SkHd3_rep_” for repeats, “SkHd3_tRNA_” for tRNAs, “SkHd3_rRNA_” for rRNAs and “SkHd3_ncRNA_” for other non-coding RNAs. These prefixes ensure global uniqueness of feature identifiers in cases where multiple annotation sources assign the same feature type (e.g., *gene* or *exon*) and allow for simple filtering of annotations by category. For example, during scRNA-seq quality control, mitochondrial genes can be easily identified using the “SkHd3_MT_” prefix to calculate the total mitochondrial gene expression per cell.

### Code Availability

No custom software was generated in the study. The scripts that we used to run the existing software are available at GitHub (https://github.com/VasiliyZubarev/HalisarcaGenome) and at Zenodo^95^ (https://doi.org/10.5281/zenodo.15992695)

## Acknowledgements

The research was done using the equipment of the Core Centrum of the Institute of Developmental Biology of the Russian Academy of Sciences and Genomics Core Facility of the Skolkovo Institute of Science and Technology. Preparation of metaphase chromosomes and genome annotation were supported by a grant from the Russian Science Foundation, number 25-14-00131 (to YL).

## Author contributions

Vasiliy Zubarev - Formal analysis; Visualisation; Data curation; Validation; Project administration; Writing - original draft; Writing - review & editing. Alexander Cherkasov - Methodology; Formal analysis; Data curation; Writing - original draft. Leonid Sidorov - Formal analysis; Visualisation; Data curation, Writing - original draft; Writing - review & editing. Kim I. Adameyko - Formal analysis; Data curation; Writing - original draft; Writing - review & editing. Alexander D. Finoshin - Investigation; Funding acquisition. Alina Ryabova - Methodology. Anastasiia Kashtanova - Visualisation. Guzel Gazizova - Investigation. Natalia Gogoleva - Investigation; Methodology. Anton V. Burakov - Investigation. Olga Kozlova - Formal analysis. Elena Shagimardanova - Investigation; Resources. Oleg Gusev - Investigation; Resources. Kirill Mikhailov - Formal analysis. Oksana Kravchuk - Conceptualization; Investigation; Supervision; Funding acquisition. Yulia Lyupina - Conceptualization; Investigation; Supervision; Funding acquisition. Ekaterina Khrameeva - Conceptualization; Supervision; Visualisation; Project administration; Writing - original draft; Writing - review & editing.

## References

1. Ereskovsky, A. V., Lavrov, D. V., Boury-Esnault, N. & Vacelet, J. Molecular and morphological description of a new species of Halisarca (Demospongiae: Halisarcida) from Mediterranean Sea and a redescription of the type species Halisarca dujardini. Zootaxa 2768, (2011).

2. Lavrov, A. I. & Kosevich, I. A. Sponge cell reaggregation: Cellular structure and morphogenetic potencies of multicellular aggregates: SPONGE CELL REAGGREGATION: CELLULAR STRUCTURE. J. Exp. Zool. Part Ecol. Genet. Physiol. 325, 158–177 (2016).

3. Ereskovsky, A. V. et al. Transdifferentiation and mesenchymal-to-epithelial transition during regeneration in Demospongiae (Porifera). J. Exp. Zoolog. B Mol. Dev. Evol. 334, 37–58 (2020).

4. Ereskovsky, A., Borisenko, I. E., Bolshakov, F. V. & Lavrov, A. I. Whole-Body Regeneration in Sponges: Diversity, Fine Mechanisms, and Future Prospects. Genes 12, 506 (2021).

5. Borisenko, I., Adamski, M., Ereskovsky, A. & Adamska, M. Surprisingly rich repertoire of Wnt genes in the demosponge Halisarca dujardini. BMC Evol. Biol. 16, 123 (2016).

6. Borisenko, I., Bolshakov, F. V., Ereskovsky, A. & Lavrov, A. I. Expression of Wnt and TGF-Beta Pathway Components during Whole-Body Regeneration from Cell Aggregates in Demosponge Halisarca dujardinii. Genes 12, 944 (2021).

7. Lavrov, A. I. et al. Cell proliferation and cell death during whole-body regeneration in the demosponge Halisarca dujardinii. FEBS Lett. 1873-3468.70025 (2025) doi:10.1002/1873-3468.70025.

8. Lyupina, Y. V. et al. The divergent intron-containing actin in sponge morphogenetic processes. NAR Genomics Bioinforma. 7, (2025).

9. Adameyko, K. I. et al. Conservative and Atypical Ferritins of Sponges. Int. J. Mol. Sci. 22, 8635 (2021).

10. Finoshin, A. D. et al. Iron metabolic pathways in the processes of sponge plasticity. PLOS ONE 15, e0228722 (2020).

11. Kravchuk, O. I. et al. Characteristics of δ-Aminolevulinic Acid Dehydratase of the Cold-Water Sponge Halisarca dujardinii. Mol. Biol. 57, 1085–1096 (2023).

12. Adameyko, K. I. et al. Structure of Neuroglobin from Cold-Water Sponge Halisarca dujardinii. Mol. Biol. 54, 416–420 (2020).

13. Finoshin, A. D. et al. Structure and Function of the Transglutaminase Cluster in the Basal Metazoan Halisarca dujardinii (Sponge). Mol. Biol. 58, 920–934 (2024).

14. Simion, P. et al. A Large and Consistent Phylogenomic Dataset Supports Sponges as the Sister Group to All Other Animals. Curr. Biol. 27, 958–967 (2017).

15. Schultz, D. T. et al. Ancient gene linkages support ctenophores as sister to other animals. Nature 618, 110–117 (2023).

16. Bethesda (MD): National Library of Medicine (US), National Center for Biotechnology Information. Genome. https://www.ncbi.nlm.nih.gov/datasets/genome/ (2004).

17. de Voogd, N. et al. World Porifera Database. Accessed at https://www.marinespecies.org/porifera. 10.14284/359 (2024).

18. Multi-omics study of the reaggregation process in the sponge Halisarca dujardinii. NCBI Sequence Read Archive https://identifiers.org/ncbi/insdc.sra:SRP519701 (2025).

19. Hybrid assembly and annotation of Halisarca dujardinii (Porifera, Demospongiae) genome. NCBI Sequence Read Archive https://identifiers.org/ncbi/insdc.sra:SRP479657 (2024).

20. Transcriptome analysis of cold water sponge Halisarca dujardini. NCBI Sequence Read Archive https://identifiers.org/ncbi/insdc.sra:SRP235047 (2019).

21. RNA-seq of Halicarca dujardinii development. NCBI Sequence Read Archive https://identifiers.org/ncbi/insdc.sra:SRP485380 (2025).

22. Tissue homeostasis at the root of Metazoa: transcriptional control of whole-body regeneration in sponges. NCBI Sequence Read Archive https://identifiers.org/ncbi/insdc.sra:SRP558447 (2025).

23. Demosponge Halisarca dujardini - transcriptome sequencing. NCBI Sequence Read Archive https://identifiers.org/ncbi/insdc.sra:ERP013253 (2016).

24. Pinaev, A. G. et al. RIAM: A Universal Accessible Protocol for the Isolation of High Purity DNA from Various Soils and Other Humic Substances. Methods Protoc. 5, 99 (2022).

25. Chen, S., Zhou, Y., Chen, Y. & Gu, J. fastp: an ultra-fast all-in-one FASTQ preprocessor. Bioinformatics 34, i884–i890 (2018).

26. Ruan, J. & Li, H. Fast and accurate long-read assembly with wtdbg2. Nat. Methods 17, 155–158 (2020).

27. Kolmogorov, M., Yuan, J., Lin, Y. & Pevzner, P. A. Assembly of long, error-prone reads using repeat graphs. Nat. Biotechnol. 37, 540–546 (2019).

28. Cheng, H., Concepcion, G. T., Feng, X., Zhang, H. & Li, H. Haplotype-resolved de novo assembly using phased assembly graphs with hifiasm. Nat. Methods 18, 170–175 (2021).

29. Hu, J. et al. NextDenovo: an efficient error correction and accurate assembly tool for noisy long reads. Genome Biol. 25, 107 (2024).

30. Sun, J., Li, R., Chen, C., Sigwart, J. D. & Kocot, K. M. Benchmarking Oxford Nanopore read assemblers for high-quality molluscan genomes. Philos. Trans. R. Soc. B Biol. Sci. 376, 20200160 (2021).

31. Shen, W., Le, S., Li, Y. & Hu, F. SeqKit: A Cross-Platform and Ultrafast Toolkit for FASTA/Q File Manipulation. PLOS ONE 11, e0163962 (2016).

32. Huang, N. & Li, H. compleasm: a faster and more accurate reimplementation of BUSCO. Bioinformatics 39, btad595 (2023).

33. Tegenfeldt, F. et al. OrthoDB and BUSCO update: annotation of orthologs with wider sampling of genomes. Nucleic Acids Res. 53, D516–D522 (2025).

34. Walker, B. J. et al. Pilon: An Integrated Tool for Comprehensive Microbial Variant Detection and Genome Assembly Improvement. PLoS ONE 9, e112963 (2014).

35. Li, H. & Durbin, R. Fast and accurate long-read alignment with Burrows–Wheeler transform. Bioinformatics 26, 589–595 (2010).

36. Guan, D. et al. Identifying and removing haplotypic duplication in primary genome assemblies. Bioinformatics 36, 2896–2898 (2020).

37. Hu, J., Fan, J., Sun, Z. & Liu, S. NextPolish: a fast and efficient genome polishing tool for long-read assembly. Bioinformatics 36, 2253–2255 (2020).

38. Santini, S. et al. The compact genome of the sponge Oopsacas minuta (Hexactinellida) is lacking key metazoan core genes. BMC Biol. 21, 139 (2023).

39. Kenny, N. J. et al. Tracing animal genomic evolution with the chromosomal-level assembly of the freshwater sponge Ephydatia muelleri. Nat. Commun. 11, 3676 (2020).

40. Wood, D. E., Lu, J. & Langmead, B. Improved metagenomic analysis with Kraken 2. Genome Biol. 20, 257 (2019).

41. Astashyn, A. et al. Rapid and sensitive detection of genome contamination at scale with FCS-GX. Genome Biol. 25, 60 (2024).

42. Dudchenko, O. et al. De novo assembly of the Aedes aegypti genome using Hi-C yields chromosome-length scaffolds. Science 356, 92–95 (2017).

43. Goloborodko, A. et al. open2c/distiller-nf: v0.3.4. Zenodo 10.5281/ZENODO.7309110 (2022).

44. Smit, A. & Hubley, R. RepeatMasker.

45. Srivastava, M., Simakov, O., Chapman, J., Fahey, B. & Gauthier, M. Genome assembly v1.1 (Amphimedon queenslandica). RefSeq https://identifiers.org/refseq.gcf:GCF_000090795.2 (2010).

46. Wellcome Sanger Tree of Life Programme. Genome assembly ooCorCand1.1 (Corticium candelabrum). RefSeq http://identifiers.org/refseq.gcf:GCF_963422355.1 (2023).

47. Wellcome Sanger Tree of Life Programme. Genome assembly odDysAvar1.4 (Dysidea avara). RefSeq http://identifiers.org/refseq.gcf:GCF_963678975.1 (2024).

48. Wellcome Sanger Tree of Life Programme. Genome assembly odHalPani1.1 (Halichondria panicea). RefSeq http://identifiers.org/refseq.gcf:GCF_963675165.1 (2023).

49. Wellcome Sanger Tree of Life Programme. Genome assembly ooOscLobu1.1 (Oscarella lobularis). RefSeq http://identifiers.org/refseq.gcf:GCF_947507565.1 (2022).

50. Wellcome Sanger Tree of Life Programme. Genome assembly ocSycCili1.1 (Sycon ciliatum). RefSeq http://identifiers.org/refseq.gcf:GCF_964019385.1 (2024).

51. Qian, J. et al. TEtrimmer: a novel tool to automate the manual curation of transposable elements. Preprint at 10.1101/2024.06.27.600963 (2024).

52. Bickmann, L., Rodriguez, M., Jiang, X. & Makałowski, W. Transformer-Based Classification of Transposable Element Consensus Sequences with TEclass2. Biology 15, 59 (2025).

53. Kojima, K. K. & Jurka, J. Crypton transposons: identification of new diverse families and ancient domestication events. Mob. DNA 2, 12 (2011).

54. Babraham Institute Bioinformatics Group. FastQC, version 0.12.0. https://www.bioinformatics.babraham.ac.uk/projects/fastqc/ (2023).

55. Ewels, P., Magnusson, M., Lundin, S. & Käller, M. MultiQC: summarize analysis results for multiple tools and samples in a single report. Bioinformatics 32, 3047–3048 (2016).

56. Grabherr, M. G. et al. Full-length transcriptome assembly from RNA-Seq data without a reference genome. Nat. Biotechnol. 29, 644–652 (2011).

57. Li, W. & Godzik, A. Cd-hit: a fast program for clustering and comparing large sets of protein or nucleotide sequences. Bioinformatics 22, 1658–1659 (2006).

58. Adamski, M. TSA: Halisarca dujardinii, transcriptome shotgun assembly. GenBank https://identifiers.org/ncbi/insdc:HADA01000000 (2019).

59. Simão, F. A., Waterhouse, R. M., Ioannidis, P., Kriventseva, E. V. & Zdobnov, E. M. BUSCO: assessing genome assembly and annotation completeness with single-copy orthologs. Bioinformatics 31, 3210–3212 (2015).

60. Hoff, K. J., Lange, S., Lomsadze, A., Borodovsky, M. & Stanke, M. BRAKER1: Unsupervised RNA-Seq-Based Genome Annotation with GeneMark-ET and AUGUSTUS. Bioinformatics 32, 767–769 (2016).

61. Brůna, T., Hoff, K. J., Lomsadze, A., Stanke, M. & Borodovsky, M. BRAKER2: automatic eukaryotic genome annotation with GeneMark-EP+ and AUGUSTUS supported by a protein database. NAR Genomics Bioinforma. 3, lqaa108 (2021).

62. Li, H. Protein-to-genome alignment with miniprot. Bioinformatics 39, btad014 (2023).

63. Brůna, T., Gabriel, L. & Hoff, K. J. Navigating Eukaryotic Genome Annotation Pipelines: A Route Map to BRAKER, Galba, and TSEBRA. Preprint at 10.48550/arXiv.2403.19416 (2024).

64. Bruna, T., Lomsadze, A. & Borodovsky, M. A new gene finding tool GeneMark-ETP significantly improves the accuracy of automatic annotation of large eukaryotic genomes. Preprint at 10.1101/2023.01.13.524024 (2023).

65. Buchfink, B., Xie, C. & Huson, D. H. Fast and sensitive protein alignment using DIAMOND. Nat. Methods 12, 59–60 (2015).

66. Gotoh, O. A space-efficient and accurate method for mapping and aligning cDNA sequences onto genomic sequence. Nucleic Acids Res. 36, 2630–2638 (2008).

67. Iwata, H. & Gotoh, O. Benchmarking spliced alignment programs including Spaln2, an extended version of Spaln that incorporates additional species-specific features. Nucleic Acids Res. 40, e161–e161 (2012).

68. Kovaka, S. et al. Transcriptome assembly from long-read RNA-seq alignments with StringTie2. Genome Biol. 20, 278 (2019).

69. Pertea, G. & Pertea, M. GFF Utilities: GffRead and GffCompare. F1000Research 9, 304 (2020).

70. Gabriel, L. et al. BRAKER3: Fully automated genome annotation using RNA-seq and protein evidence with GeneMark-ETP, AUGUSTUS, and TSEBRA. Genome Res. 34, 769–777 (2024).

71. Stanke, M., Diekhans, M., Baertsch, R. & Haussler, D. Using native and syntenically mapped cDNA alignments to improve de novo gene finding. Bioinformatics 24, 637–644 (2008).

72. Stanke, M., Schöffmann, O., Morgenstern, B. & Waack, S. Gene prediction in eukaryotes with a generalized hidden Markov model that uses hints from external sources. BMC Bioinformatics 7, 62 (2006).

73. Gabriel, L., Hoff, K. J., Brůna, T., Borodovsky, M. & Stanke, M. TSEBRA: transcript selector for BRAKER. BMC Bioinformatics 22, 566 (2021).

74. Dobin, A. et al. STAR: ultrafast universal RNA-seq aligner. Bioinformatics 29, 15–21 (2013).

75. Danecek, P. et al. Twelve years of SAMtools and BCFtools. GigaScience 10, giab008 (2021).

76. Robinson, J. T. et al. Integrative genomics viewer. Nat. Biotechnol. 29, 24–26 (2011).

77. Haas, B. J. Improving the Arabidopsis genome annotation using maximal transcript alignment assemblies. Nucleic Acids Res. 31, 5654–5666 (2003).

78. Haas, B. J. et al. Automated eukaryotic gene structure annotation using EVidenceModeler and the Program to Assemble Spliced Alignments. Genome Biol. 9, R7 (2008).

79. Zheng, G. X. Y. et al. Massively parallel digital transcriptional profiling of single cells. Nat. Commun. 8, 14049 (2017).

80. Haese-Hill, W., Crouch, K. & Otto, T. D. peaks2utr: a robust Python tool for the annotation of 3′ UTRs. Bioinformatics 39, btad112 (2023).

81. Chan, P. P., Lin, B. Y., Mak, A. J. & Lowe, T. M. tRNAscan-SE 2.0: improved detection and functional classification of transfer RNA genes. Nucleic Acids Res. 49, 9077–9096 (2021).

82. Bryant, D. M. et al. A Tissue-Mapped Axolotl De Novo Transcriptome Enables Identification of Limb Regeneration Factors. Cell Rep. 18, 762–776 (2017).

83. Eddy, S. R. Accelerated Profile HMM Searches. PLoS Comput. Biol. 7, e1002195 (2011).

84. Moriya, Y., Itoh, M., Okuda, S., Yoshizawa, A. C. & Kanehisa, M. KAAS: an automatic genome annotation and pathway reconstruction server. Nucleic Acids Res. 35, W182–W185 (2007).

85. Cantalapiedra, C. P., Hernández-Plaza, A., Letunic, I., Bork, P. & Huerta-Cepas, J. eggNOG-mapper v2: Functional Annotation, Orthology Assignments, and Domain Prediction at the Metagenomic Scale. Mol. Biol. Evol. 38, 5825–5829 (2021).

86. Huerta-Cepas, J. et al. Fast Genome-Wide Functional Annotation through Orthology Assignment by eggNOG-Mapper. Mol. Biol. Evol. 34, 2115–2122 (2017).

87. Steinegger, M. & Söding, J. MMseqs2 enables sensitive protein sequence searching for the analysis of massive data sets. Nat. Biotechnol. 35, 1026–1028 (2017).

88. Teufel, F. et al. SignalP 6.0 predicts all five types of signal peptides using protein language models. Nat. Biotechnol. 40, 1023–1025 (2022).

89. Krogh, A., Larsson, B., Von Heijne, G. & Sonnhammer, E. L. L. Predicting transmembrane protein topology with a hidden markov model: application to complete genomes. J. Mol. Biol. 305, 567–580 (2001).

90. Jin, J.-J. et al. GetOrganelle: a fast and versatile toolkit for accurate de novo assembly of organelle genomes. Genome Biol. 21, 241 (2020).

91. Fiedler, L., Bernt, M. & Middendorf, M. DeGeCI 1.1: a web platform for gene annotation of mitochondrial genomes. Bioinforma. Adv. 4, vbae072 (2024).

92. Donath, A. et al. Improved annotation of protein-coding genes boundaries in metazoan mitochondrial genomes. Nucleic Acids Res. 47, 10543–10552 (2019).

93. Grant, J. R. et al. Proksee: in-depth characterization and visualization of bacterial genomes. Nucleic Acids Res. 51, W484–W492 (2023).

94. Zubarev, V., Cherkasov, A., Sidorov, L. & Adameyko, K. Genome assembly SKOL_HDuj_3.0 GenBank https://identifiers.org/insdc.gca:GCA_054858975 (2026).

95. Zubarev, V. et al. Chromosome-level genome assembly of the sponge Halisarca dujardinii. Zenodo 10.5281/ZENODO.15992694 (2026).

96. Jacques Dainat et al. NBISweden/AGAT: AGAT-v1.4.1. Zenodo 10.5281/ZENODO.3552717 (2024).

97. Marçais, G. & Kingsford, C. A fast, lock-free approach for efficient parallel counting of occurrences of k-mers. Bioinformatics 27, 764–770 (2011).

98. Vurture, G. W. et al. GenomeScope: fast reference-free genome profiling from short reads. Bioinformatics 33, 2202–2204 (2017).

99. Ranallo-Benavidez, T. R., Jaron, K. S. & Schatz, M. C. GenomeScope 2.0 and Smudgeplot for reference-free profiling of polyploid genomes. Nat. Commun. 11, 1432 (2020).

100. Borisenko, I., Predeus, A., Lavrov, A. & Ereskovsky, A. First draft genome assembly of sponge Halisarca dujardinii reveals key components of basement membrane and broad repertoire of aggregation factors. Sci. Rep. 15, 44778 (2025).

101. Borisenko, I., Predeus, A., Lavrov, A. & Ereskovsky, A. Genome assembly ASM4999745v1. GenBank https://identifiers.org/insdc.gca:GCA_049997455.1 (2024).

102. Dynamic changes in gene expression in regeneration of Halisarca dujardinii (Porifera, Demospongiae). NCBI Sequence Read Archive https://identifiers.org/ncbi/insdc.sra:SRP485609 (2024).

